# Timing of fruiting rather than topography determines the direction of vertical seed dispersal by mammals and birds

**DOI:** 10.1101/2020.05.24.112987

**Authors:** Y. Tsunamoto, S. Koike, I. Tayasu, T. Masaki, S. Kato, S. Kikuchi, T. Nagamitsu, T. Haraguchi, T. Naganuma, S. Naoe

## Abstract

Vertical seed dispersal toward higher or lower altitudes has been recognized as one of the critical processes for plants to escape from climate change. Studies exploring vertical seed dispersal are scarce, preventing the prediction of future vegetation dynamics. In the present study, we show that the timing of fruiting, rather than topography, determines the direction of vertical seed dispersal by mammals and birds across mountains in central Japan. We found strong uphill seed dispersal of summer fruiting cherry and weak downhill seed dispersal of summer-to-autumn fruiting cherry, irrespective of mountains and animals. The ascent or descent of animals, following the altitudinal gradients in food plant phenology in the temperate zone, was considered to be a driver of the biased seed dispersal. We found that megafauna (i.e., bears) intensively dispersed seeds vertically. The results suggest that the timing of fruiting and megafauna strongly affect whether animal-dispersed temperate plants can maintain their populations under climate change.

## INTRODUCTION

Vertical seed dispersal, that is, seed dispersal toward higher or lower altitudes, has now been recognized as one of the critical processes for plant migration, especially under climate change (*1, 2*). Uphill or downhill seed dispersal is advantageous or disadvantageous, respectively, for plant escape and expansion under global warming, and vice versa under global cooling (*3*). Under current global warming, which shifts suitable plant habitats to colder areas (i.e., higher altitudes and latitudes) (*4*), uphill seed dispersal is essential for plants to maintain their populations. Seed dispersal toward higher latitudes is also beneficial for plants; however, vertical seed dispersal is far more efficient than latitudinal one because the temperature decrease with increasing altitude is 100–1000 times more than that of an equivalent latitudinal distance (*5*).^1^

The prediction of vertical seed dispersal by animals is considered to be difficult compared to other seed dispersal modes that utilize abiotic physical factors, such as wind and water flow. Many previous studies on horizontal seed dispersal have shown that animal behaviors that determine seed dispersal patterns are affected by various factors at different spatial scales (*6–8*). The fact that the seeds of one plant species are generally dispersed by various animal species (*9*) makes the seed dispersal pattern more complex. In vertical seed dispersal, altitudinal changes in biotic and abiotic factors, including food resources and climates, are also likely to affect the behavior of frugivores (i.e., fruit-eating animals). The difficulty in predicting vertical seed dispersal by animals largely prevents the prediction of future forest dynamics under global warming because many of trees, in terms of the number of species and abundance, depend on animals for seed dispersal (*10*). At present, it is important to accumulate observations of vertical seed dispersal in the field and set up hypotheses based on them. However, there are only three studies that have investigated vertical seed dispersal itself, due to the technical difficulty or high cost (*1–3*).

Recently, Naoe et al. hypothesized that the timing of fruiting determines the direction of vertical seed dispersal: uphill seed dispersal occurs in spring- and summer-fruiting plants and downhill seed dispersal occurs in autumn- and winter-fruiting plants because frugivores ascend or descend mountains following food plant phenology in the temperate zone (*1*), which proceeds from the foot to the top of mountains in spring–summer and from the top to the foot in autumn– winter (*11, 12*). Considering that most animal-dispersed trees fruit in autumn–winter in the temperate zone (*10*), this hypothesis suggests that populations of autumn-to-winter fruiting trees dispersed by animals may not sufficiently escape from current global warming in the temperate zone (*3*). However, this hypothesis is based on observations of vertical seed dispersal by mammals on one mountain range, and thus its generality should be tested in several aspects, including topography, the extinction of megafauna, and seed dispersal by birds. Previous studies did not explicitly evaluate the effect of topography. On steep mountains, farther vertical seed dispersal is expected to occur compared to gently sloping mountains because frugivores can ascend and descend more efficiently (*2, 3*). Surface areas at lower altitudes are generally larger than those at higher altitudes (*13*), and thus downhill seed dispersal is likely to occur if frugivores move randomly in the mountains (*3*). Human activities represented by habitat fragmentation and hunting can dramatically change frugivore composition, often resulting in the local extinction of megafauna, which plays an important role in seed dispersal (*14–16*). Therefore, vertical seed dispersal patterns can differ depending on the megafauna that are present/extinct in the mountains. Considering that these aspects can vary among mountains, it is essential to examine whether this hypothesis is consistently supported irrespective of the mountains. In addition, it is also essential to examine whether this hypothesis is supported in vertical seed dispersal by birds, which are another major group of frugivores (*10*).

In this study, our study aim was to clarify whether the timing of fruiting determines the direction of vertical seed dispersal by mammals and birds, irrespective of the mountains. We set up three specific questions: 1) Does the timing of fruiting and/or topography affect the direction of vertical seed dispersal? 2) Do patterns of vertical seed dispersal differ between by mammals and birds? and 3) Does the extinction of megafauna limit vertical seed dispersal? To answer these questions, we investigated the vertical seed dispersal of two cherry species (*Cerasus leveilleana* and *Padus grayana*) by mammals and birds in the Ashio-Nikko Mountains (hereafter, ASH), Abukuma Highlands (ABU), and Kanto Mountains (KAN) in central Japan (**Fig. 1AB**; **Table S1**). The two cheery species have similar fruit traits (e.g., fruit size and nutrition) and share frugivores; however, they fruit at different times (*17*), providing a good opportunity to evaluate whether the timing of fruiting affects vertical seed dispersal. We estimated the vertical seed dispersal distance by using the altitudinal gradient of the oxygen isotope ratios of seeds to locate the altitudes of the mother trees (*18*) (**Fig. S1**). The fauna of frugivores are relatively well-preserved in three mountains; however, frugivorous megafauna in Japan, that is, the Asian black bear (*Ursus thibetanus*), has become locally extinct in ABU. In ABU, Japanese macaque (*Macaca fuscata*) has also become locally extinct. As for KAN, we reanalyzed the data of a study (*1*) that reported on vertical seed dispersal by mammals. To examine the effect of topography on the direction of vertical seed dispersal, we estimated the expected vertical seed dispersal by simulation, assuming that frugivores randomly move and disperse seeds based on the topography in their home ranges, and compared the expected and observed vertical seed dispersals.

**Fig. 1.**
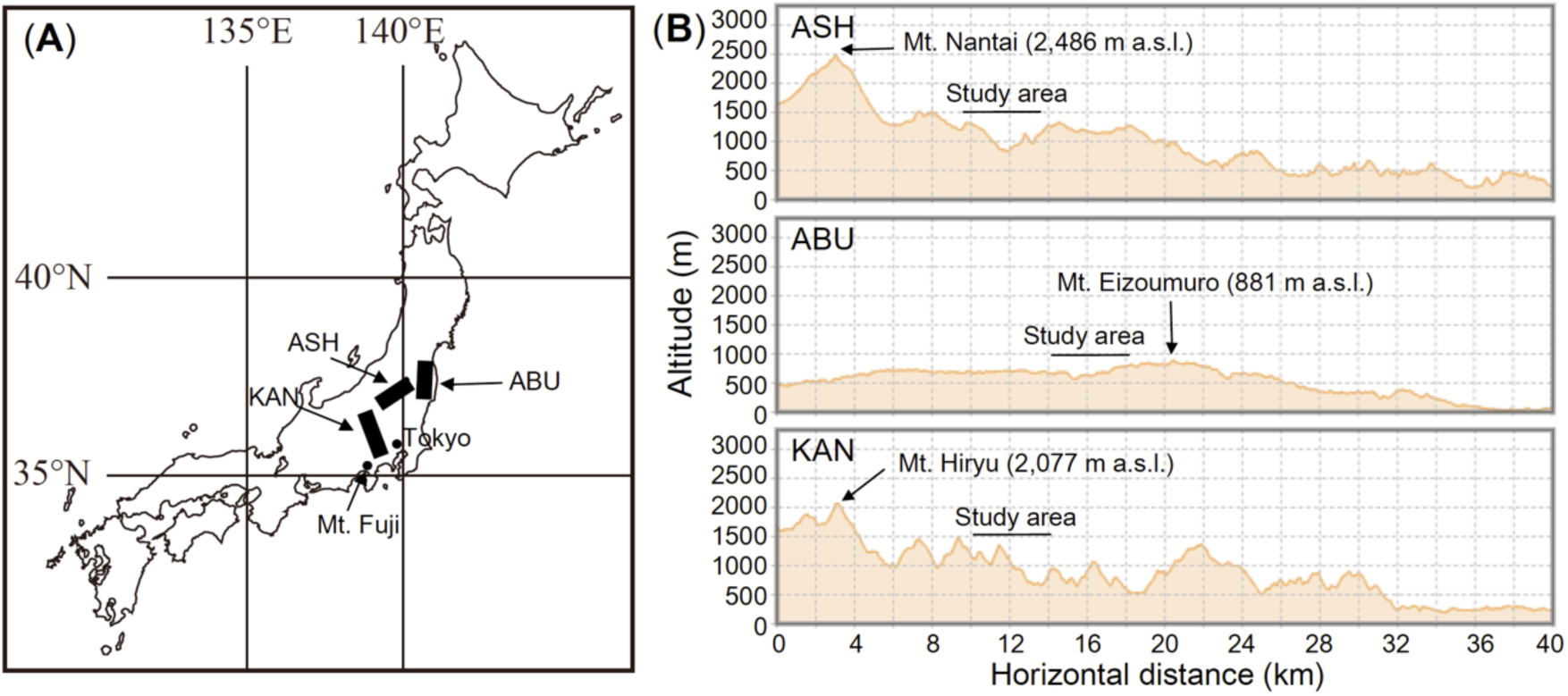
Locations of the Ashio-Nikko Mountains (ASH), Abukuma Highlands (ABU), Kanto Mountains (KAN) (**A**), and the rough topography around the study areas (**B**). Topography was obtained from the Geospatial Information Authority of Japan (https://maps.gsi.go.jp/).

## RESULTS

### Amount of seed dispersal

The number of mammal-dispersed *Cerasus leveilleana* seeds in ASH was much higher than that in ABU, where bears and macaques are locally extinct, in both study years; 1,937 vs. 724 seeds in 2014, and 5,147 vs. 513 in 2015, respectively (**Fig. 2A**; **Table S2**). In ASH, bears, macaques, martens, and raccoon dogs accounted for 43.9%, 42.0%, 9.0%, and 5.1% of the dispersed seeds in 2014, and 58.8%, 15.0%, 26.2%, and 0.0% in 2015, respectively. In ABU, martens and raccoon dogs accounted for 79.7 and 20.3% in 2014, and 100.0% and 0.0% in 2015, respectively. The number of bird-dispersed *C. leveilleana* seeds in ASH was greater than that in ABU in both study years; 29 vs. 8 seeds in 2014, and 105 vs. 14 in 2015, respectively (**Fig. 2B**; **Table S2**).

**Fig. 2.**
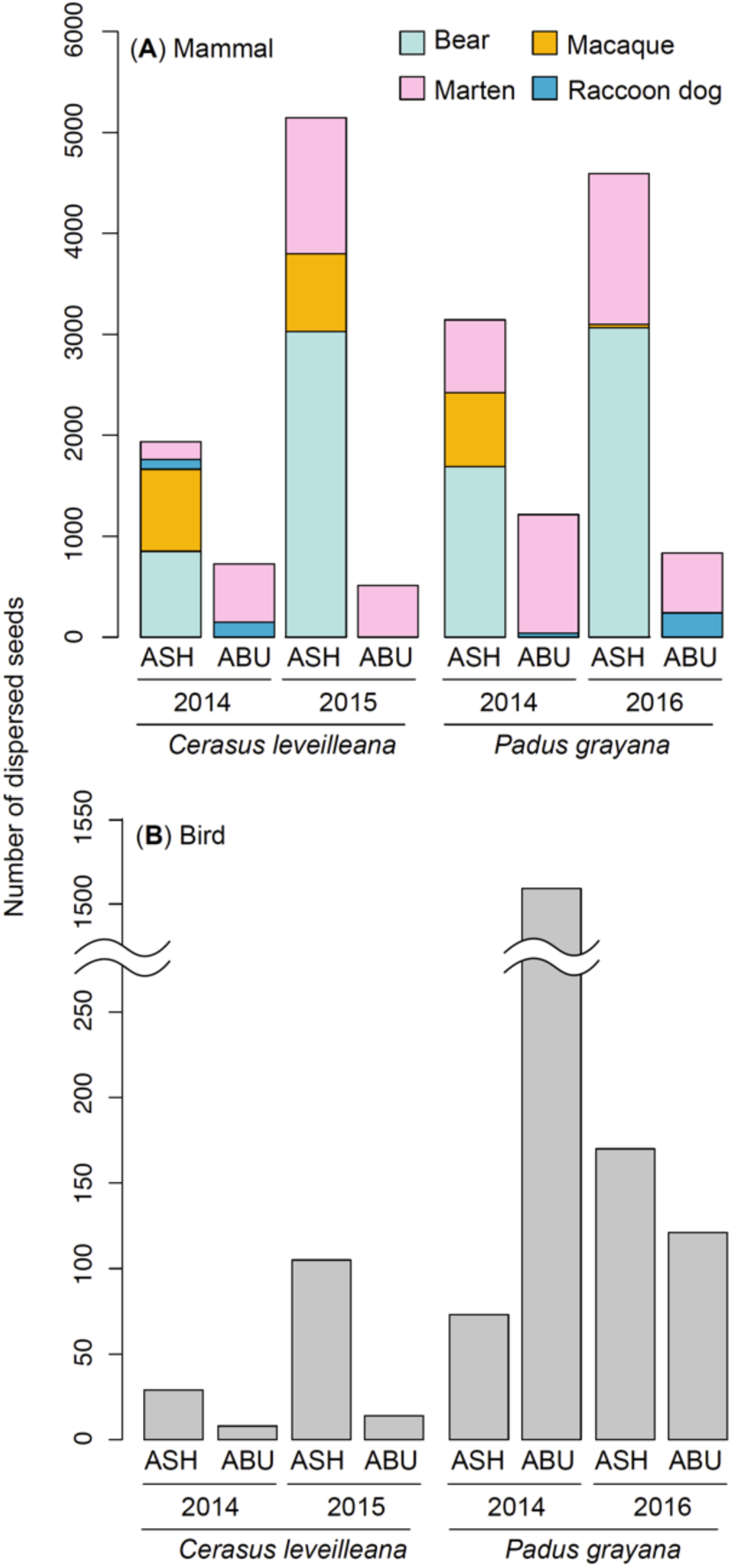
Number of dispersed seeds of *Cerasus leveilleana* and *Padus grayana* by (**A**) each mammal species and (**B**) bird.

The number of mammal-dispersed *Padus grayana* seeds in ASH was much higher than that in ABU in both study years; 3,144 vs. 1,215 seeds in 2014, and 4,593 vs. 835 in 2016 (**Fig. 2A**; **Table S2**). In ASH, bears, macaques, martens, and raccoon dogs accounted for 53.7%, 23.3%, 23.0%, and 0.0% of the dispersed seeds in 2014, and 66.8%, 0.7%, 32.5%, and 0.7% of dispersed seeds in 2015, respectively. In ABU, martens and raccoon dogs accounted for 96.7% and 3.3% of the dispersed seeds in 2014, and 71.3% and 28.7% of dispersed seeds in 2015, respectively. The number of bird-dispersed *C. leveilleana* seeds in ASH was much lower than that in ABU in 2014, but the number in ASH was greater than that in ABU in 2016; 73 vs. 1,509 seeds in 2014, and 170 vs. 121 in 2016, respectively (**Fig. 2B**; **Table S2**).

### Calibration lines for the estimation of vertical seed dispersal distances

Calibration lines for the vertical seed dispersal estimates were successfully constructed for both *C. leveilleana* and *P. grayana*; negative correlations between the altitudes and isotope ratio of non-dispersed reference seeds were detected for all sites and species (**Fig. S1AB**; **Table S3**). The *R^* values of all the parameters in the Bayesian inference for estimating calibration lines were <1.01, indicating that our model convergence was good (*19*). The ranges of isotope ratios of non-dispersed *C. leveilleana* and *P. grayana* seeds almost fully covered those of dispersed seeds. The ranges of isotope ratios of non-dispersed and dispersed *C. leveilleana* seeds were 24.18‰– 36.25‰ vs. 28.76‰–35.42‰ in ASH, and 18.53‰–35.89‰ vs. 21.40‰–32.63‰ in KAN, respectively. The ranges of isotope ratios of non-dispersed and dispersed *P. grayana* seeds were 20.55‰–28.53‰ vs. 22.12‰–29.16‰ in ASH, and 21.88‰–29.00‰ vs. 21.47‰–26.86‰ in ABU, respectively. These results indicate that the effect of extrapolation is negligible when we estimated the vertical seed dispersal distance. The *R^* values of all the parameters in the Bayesian inference for estimating vertical seed dispersal distance were <1.06, indicating that our model convergence was good.

### Vertical seed dispersal distance

For *C. leveilleana*, which fruits in early summer, the direction of estimated seed dispersal was biased toward the top of the mountains regardless of the mountains and frugivores (**Fig. 3**; **Table 1**). In ASH, bears, macaques, martens, and birds intensively dispersed seeds uphill. In KAN, bears and martens intensively dispersed seeds uphill. Differences in the vertical seed dispersal distance among frugivores were not clear in ASH (bear: mean +292.9 ± 29.3 m SE, macaque: +314.6 ± 20.9 m, marten: +333.6 ± 18.6 m, bird: +328.8 ± 26.6 m) (**Fig. 3**), while bears dispersed seeds uphill farther compared to martens in KAN (bear: +310.7 ± 33.3 m, marten: +196.3 ± 27.3 m). The absolute vertical seed dispersal distance was nearly identical to the vertical seed dispersal distance in each species in both ASH and KAN (**Table S4**), because downhill seed dispersal rarely occurred. In the simulation, assuming that frugivores randomly move and disperse seeds in their home ranges based on the topography, the expected mean vertical seed dispersal was +3.9, +4.4, -11.3, -48.8, and 1.8 m in ASH, when *r* = 0.1, 0.5, 1, 3, and 10 km, respectively (**Fig. 4**). These results indicate that strong uphill or downhill seed dispersal rarely occurs if frugivores disperse seeds randomly. The expected mean vertical seed dispersal distance was ca. 300 m less than that observed by all frugivores in ASH. In KAN, the expected mean vertical seed dispersal was +11.8, +54.0, +113.9, +157.2, and +156.1 m, respectively. These results indicate that uphill seed dispersal occurs if frugivores disperse seeds randomly, but the values were ca. 100–150 m less than those observed by bears and martens in KAN. The expected absolute mean vertical seed dispersal distance in KAN was greater than that in ASH irrespective of the *r* values (**Table S4**), indicating that KAN is steeper than ASH, and thus a farther vertical seed dispersal distance occurs in KAN than in ASH if frugivores move randomly. However, there was no such tendency in the observed absolute vertical seed dispersal distance between KAN and ASH (**Table S4**).

**Table 1.**
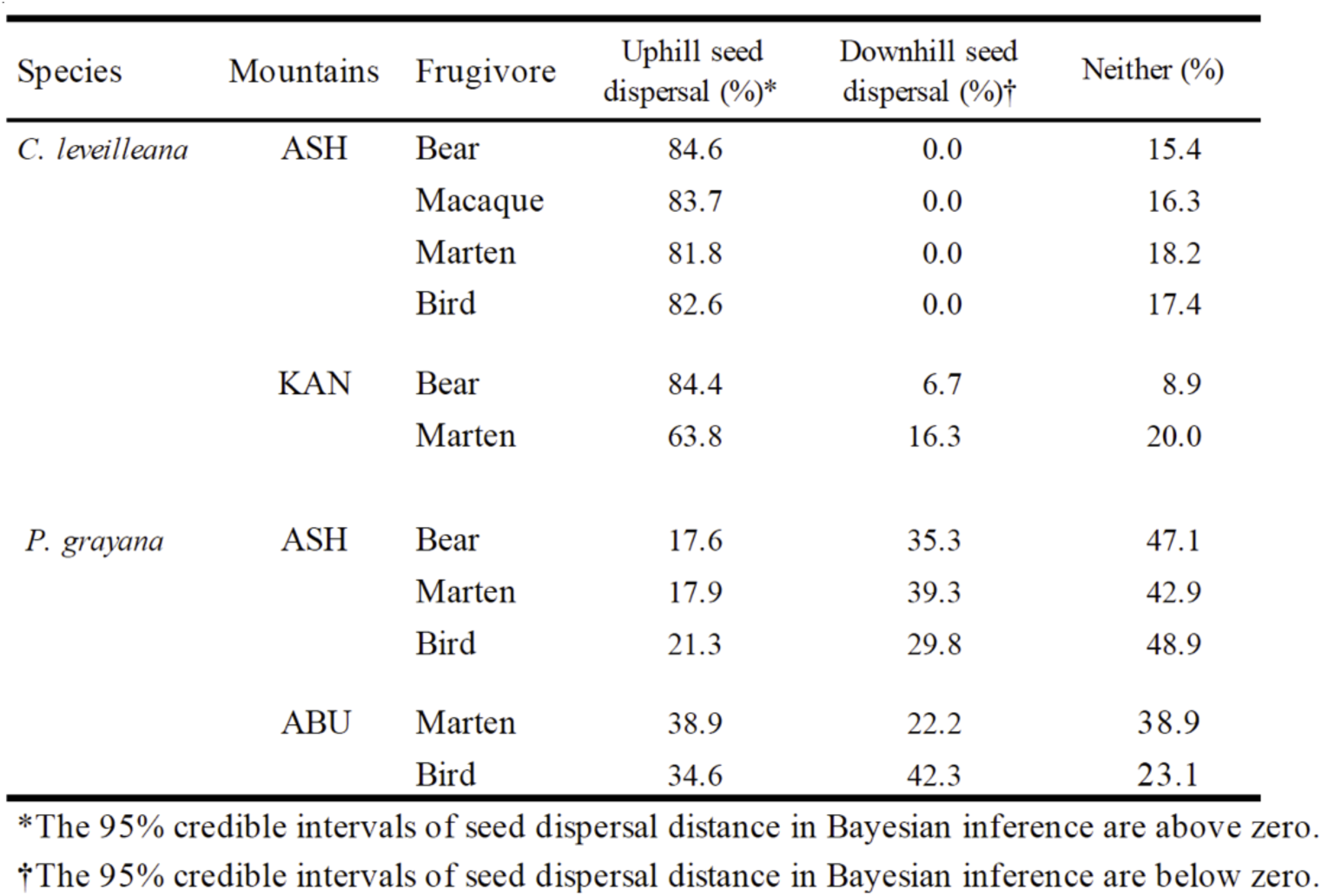
The direction of vertical seed dispersal of *Cerasus leveilleana* and *Padus grayana* in the Ashio-Nikko Mountains (ASH), Abukuma Highlands (ABU), and Kanto Mountains (KAN).

| Species | Mountains | Frugivore | Uphill seed dispersal (%)<sup>*</sup> | Downhill seed dispersal (%)<sup>†</sup> | Neither (%) |
| --- | --- | --- | --- | --- | --- |
| *C. leveilleana* | ASH | Bear | 84.6 | 0.0 | 15.4 |
|  |  | Macaque | 83.7 | 0.0 | 16.3 |
|  |  | Marten | 81.8 | 0.0 | 18.2 |
|  |  | Bird | 82.6 | 0.0 | 17.4 |
|  | KAN | Bear | 84.4 | 6.7 | 8.9 |
|  |  | Marten | 63.8 | 16.3 | 20.0 |
| *P. grayana* | ASH | Bear | 17.6 | 35.3 | 47.1 |
|  |  | Marten | 17.9 | 39.3 | 42.9 |
|  |  | Bird | 21.3 | 29.8 | 48.9 |
|  | ABU | Marten | 38.9 | 22.2 | 38.9 |
|  |  | Bird | 34.6 | 42.3 | 23.1 |

<sup>\*</sup>The 95% credible intervals of seed dispersal distance in Bayesian inference are above zero.
<sup>†</sup>The 95% credible intervals of seed dispersal distance in Bayesian inference are below zero.

**Fig. 3.**
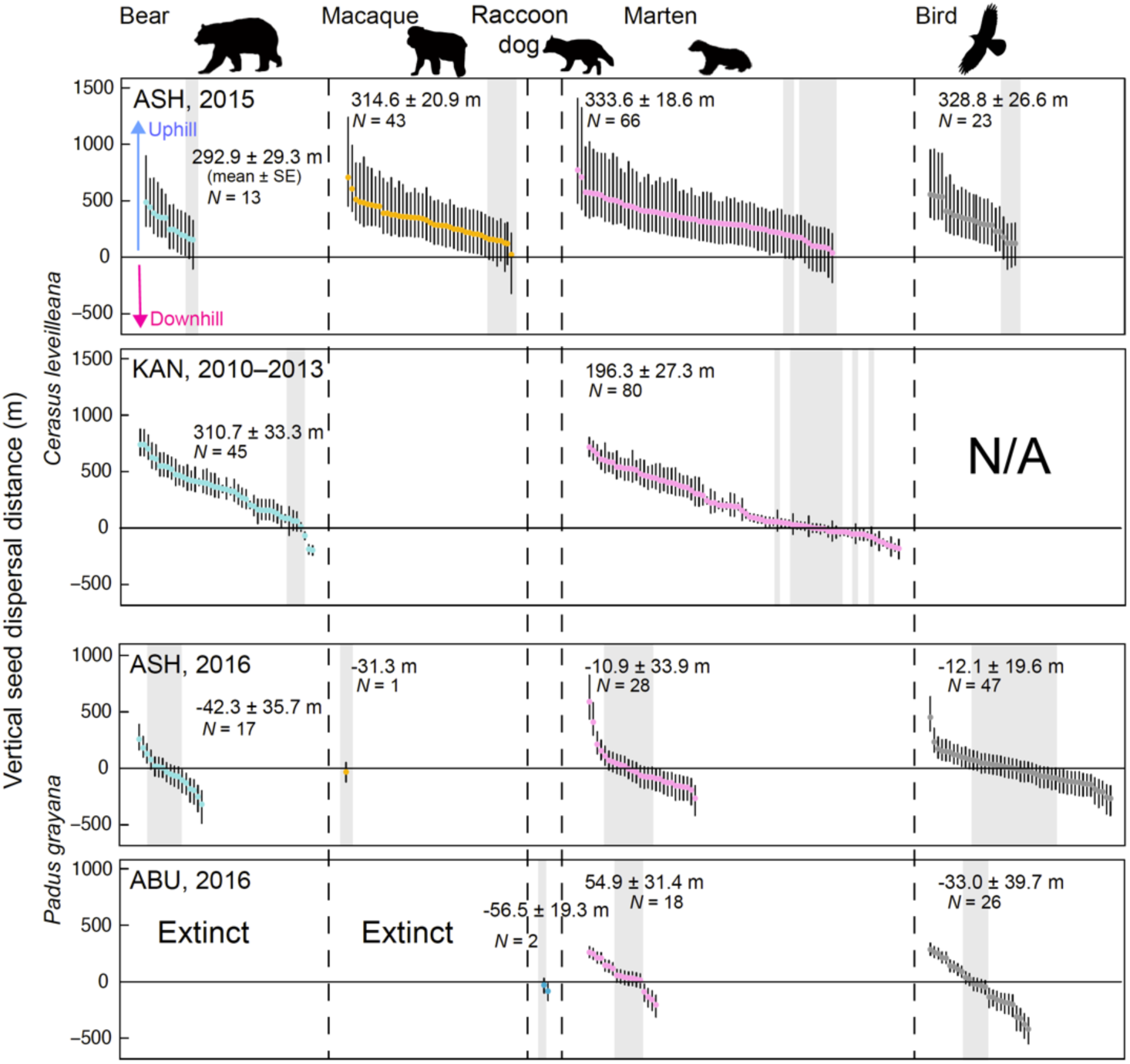
The estimated vertical seed dispersal distance by each mammal species and bird in each mountains. The dots indicate the mean vertical distance, and the vertical bars indicate the 95% credible interval of each dispersed seed. The seeds with a 95% credible interval crossed zero are shown in a gray background. Silhouette images of the bear, marten, and bird (by L. Shyamal) were obtained from PhyloPic (http://phylopic.org).

**Fig. 4.**
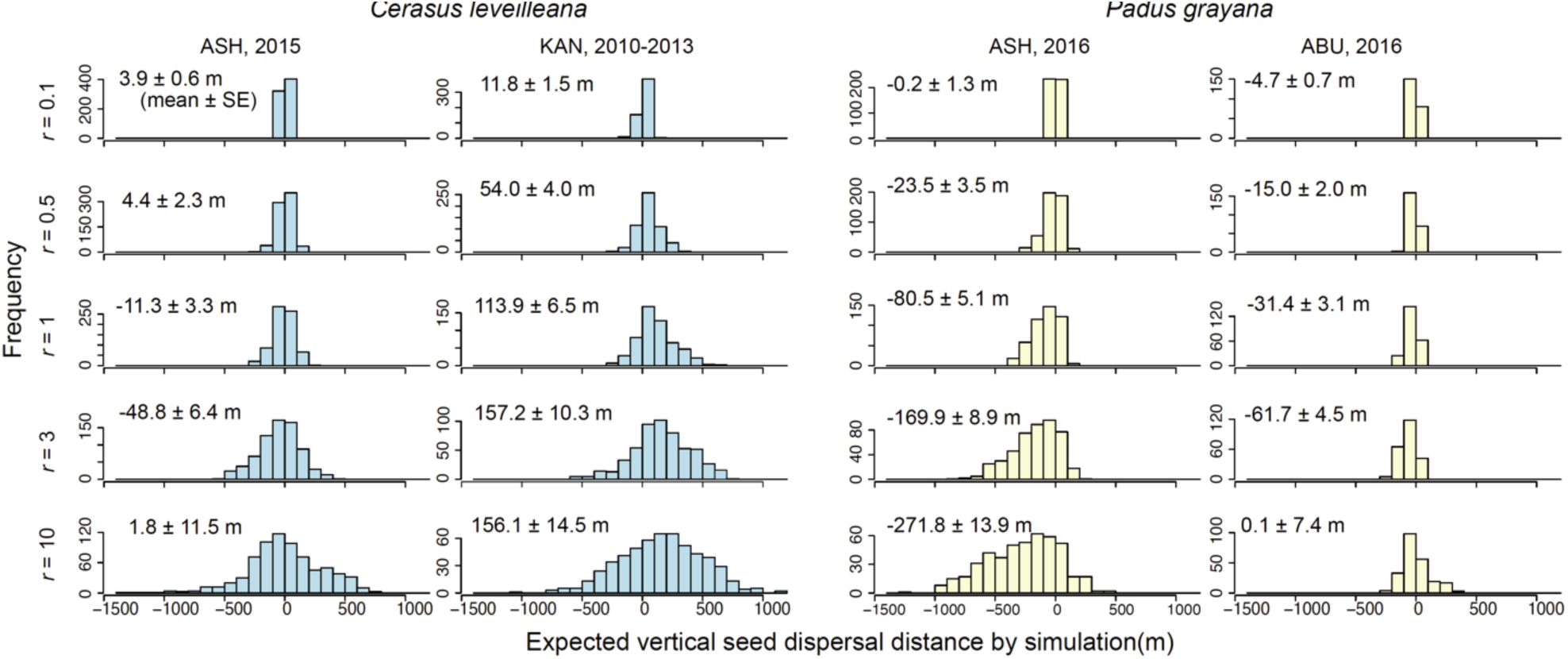
The expected vertical seed dispersal distance by simulation assuming that frugivores move and disperse seeds randomly in their home range (*r* = 0.1, 0.5, 1, 3, and 10 km). Each *r* value is roughly correspondent to the home range sizes of small birds, intermediate-sized birds, large birds and intermediate-sized mammals, Asian black bears in KAN, and Asian black bears in ASH, respectively.

For *P. grayana*, which fruits from late summer to early autumn, the direction of estimated seed dispersal was biased toward the foot of the mountains regardless of the mountains and frugivores, except for martens in ABU, but the tendency was weak (**Fig. 3**; **Table 1**). In ASH, bears, martens, and birds tended to disperse seeds downhill. In ABU, martens tended to disperse seeds uphill, and birds tended to disperse seeds downhill. Differences in the vertical seed dispersal distance among the main frugivores were not clear in ASH (bear: -42.3 ± 35.7 m, marten: -10.9 ± 33.9 m, bird: -12.1 ± 19.6 m)(**Fig. 3**), while the vertical seed dispersal distance by martens was greater than that by birds in ABU (marten: +54.9 ± 31.4 m, bird: -33.0 ± 39.7 m). Differences in the absolute vertical seed dispersal distance among frugivores were not clear in both ASH and ABU (**Table S4**). In the simulation based on topography, the expected mean vertical seed dispersal was -0.2, -23.5, -80.5, -169.9, and -271.8 m in ASH, when *r* = 0.1, 0.5, 1, 3, and 10 km, respectively (**Fig. 4**). This indicates that weak or strong downhill seed dispersal occurs if small-to-intermediate-sized frugivores or bears disperse seeds randomly. The values were relatively similar to those by all frugivores except for bears in ASH. In ABU, the expected mean vertical seed dispersal was -4.7, -15.0, -31.4, -61.7, and +0.1 m, indicating weak downhill seed dispersal. The values were relatively similar to those by birds, but not to those by martens in ABU. The expected absolute mean vertical seed dispersal distance in ASH was much greater than that in ABU irrespective of the *r* values (**Table S4**), indicating that ASH is much steeper than ABU, and thus that a farther vertical seed dispersal distance occurs in ASH compared to ABU if frugivores move randomly. However, there was no such tendency in the observed absolute vertical seed dispersal distance between KAN and ABU (**Table S4**).

### Contribution of each frugivore to whole vertical seed dispersal

When we combined the amount of seed dispersal and vertical seed dispersal distance by each mammal, the contribution of bears to the seed dispersal of *C. leveilleana* was apparent in both ASH and KAN: bears dispersed many seeds over long distances (**Fig. 5A**). Similarly, the contribution of bears to the seed dispersal of *P. grayana* was apparent in ASH (**Fig. 5B**). In ABU, where bears have become extinct, the amount of seed dispersal and long-distance vertical seed dispersal were highly limited. In ABU, birds seemed to compensate for the lack of long-distance vertical seed dispersal by bears, although we could not compare their contribution to that of mammals in terms of the amount of seed dispersal because of the different sampling methods.

**Fig. 5.**
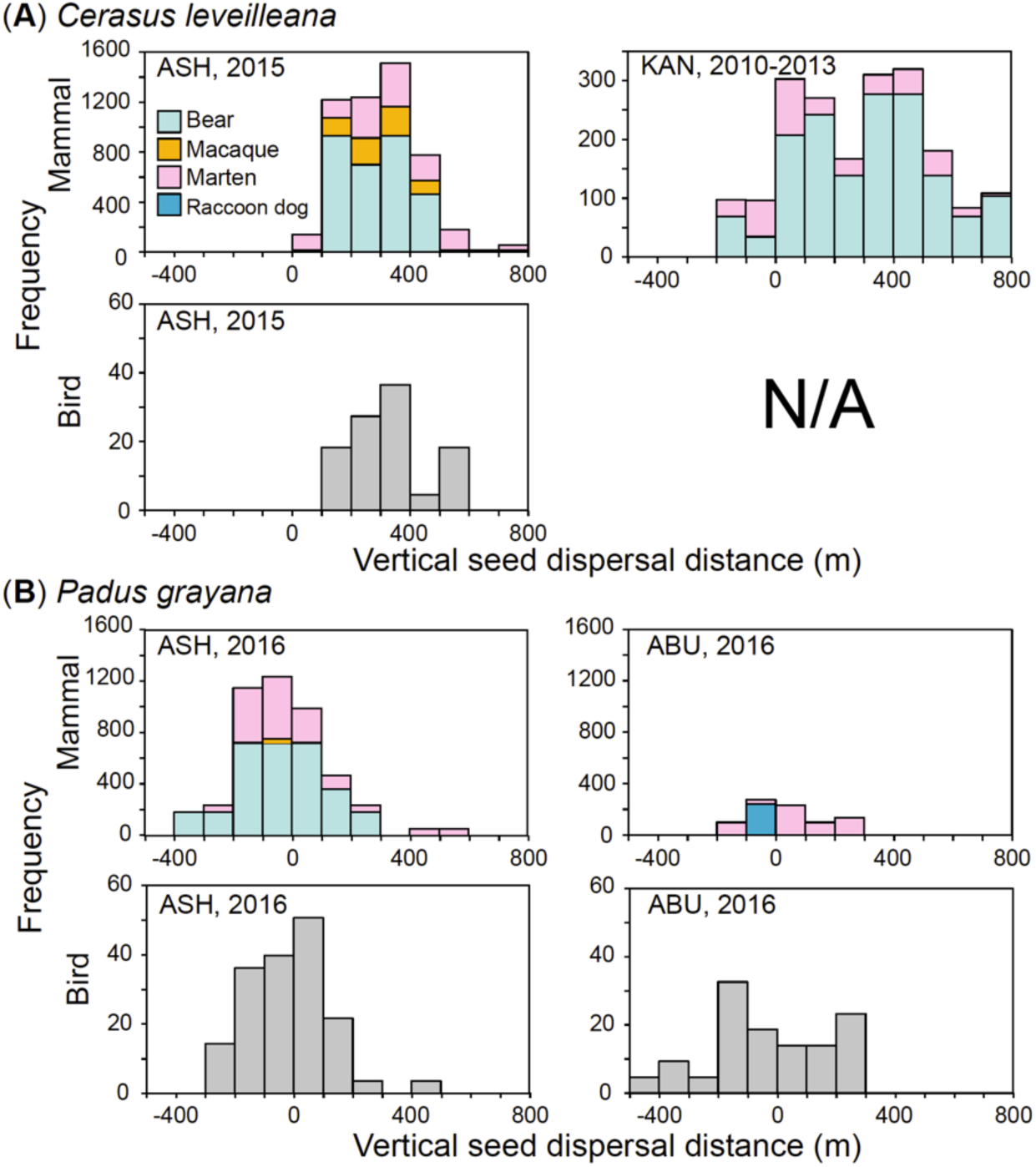
Frequency distribution of dispersed seeds of (**A**) *Cerasus leveilleana* and (**B**) *Padus grayana* by each mammal species and bird.

## DISCUSSION

The results of the two cherry species, which share frugivores but fruit at different times, supported the hypothesis that uphill seed dispersal occurs in spring- and summer-fruiting plants, and downhill seed dispersal occurs in autumn- and winter-fruiting plants owing to the movement of frugivores following food plant phenology in the temperate zone, irrespective of the mountains and frugivores. In addition to that previously reported in KAN, we observed strong uphill seed dispersal of *C. leveilleana*, which fruits in early summer, in ASH. All frugivores, that is, bears, macaques, martens, and birds, intensively dispersed seeds uphill. The frugivores’ simple tracking of ascending food plant phenology, and their daily foraging pattern—movement toward lower altitudes for cherry fruits and subsequent movement toward higher altitudes for young edible vegetation (i.e., herbs, buds, young leaves, and flowers) and accompanying larvae of herbivorous insects that are no longer abundant at lower altitudes because of the phenological progress—are considered to be the reasons for uphill seed dispersal (*1*). We observed the weak downhill seed dispersal of *P. grayana*, which fruits in late summer to early autumn in ASH and ABU. All frugivores, except martens in ABU (which dispersed seeds +54.9 m uphill), dispersed seeds downhill. Vertical seed dispersal distances of *P. grayana* (i.e., -56.5–+54.9 m) were intermediate values of those of summer-fruiting *C. leveilleana* (ca. +200–+300 m) and those of autumn-fruiting *Actinidia arguta* (ca. -300–-100 m, except for raccoon dogs) (*3*). This is probably because there were no strong altitudinal gradients in the phenology of food plants at the time of *P. grayana* fruiting, which is during the transitional period between summer and autumn. During this period, frugivores would not have a strong motivation to ascend or descend the mountains, resulting in less biased vertical seed dispersal.

Frugivores did not move and disperse seeds simply based on the topography of mountains, indicating that the effect of topography on the direction of vertical seed dispersal is relatively small compared to that of the timing of fruiting. The observed vertical seed dispersal distances of *C. leveilleana* in ASH and KAN were much greater than the expected one assuming that frugivores move and disperse seeds randomly based on the topography around the census routes. In addition, the previously reported vertical seed dispersal distance of *A. arguta* in KAN (*3*) was much less (ca. 100–450 m) than the expected one of *C. leveilleana*, which were estimated using the same census route as *A. arguta*. In KAN, which is steeper than ASH, farther vertical seed dispersal compared to ASH was expected, but such a tendency was not detected. The observed vertical seed dispersal distances of *P. grayana* in ASH and ABU, which showed weak downhill seed dispersal (except by martens in ABU), were relatively similar to the expected ones based on the topography. You might think that frugivores moved and dispersed seeds randomly when they did not have strong motivation to ascend or descend the mountains. However, the observed vertical seed dispersal distance by bears in ASH (mean of -42.3 m) was much greater than the expected one (mean of -271.8 m), which corresponds to their home range in ASH (i.e., *r* = 10 km). This indicates that bears maintained their altitudinal locations, and thus moved and dispersed seeds non-randomly at the fruiting time of *P. grayana. Padus grayana* fruits from August to mid-September, which is the hottest period in Japan. During this period, the bears in ASH decrease their daily activity (*20*) partly because they need to maintain their metabolism (Koike unpublished data). Their reluctance to move, especially toward lower and thus warmer altitudes, is likely to explain this non-random seed dispersal. In ASH, which is far steeper than ABU, farther vertical seed dispersal compared to ABU was expected, but such a tendency was not detected. Combined with the gaps between the observed and expected vertical seed dispersal of *P. grayana*, it was considered that frugivores do not simply move and disperse seeds randomly based on topography, even when there are no strong altitudinal gradients in the phenology of food plants. In this study, we expected that the surface areas at lower altitudes are larger than those at higher altitudes (*13*), resulting in downhill seed dispersal if frugivores move randomly. However, the distributions of the expected vertical seed dispersal distance by simulation were not clearly biased toward lower altitudes, even if *r* = 10 km, in three out of four cases (i.e., *C. leveilleana* in ASH, and *P. grayana* in ASH and ABU) (**Fig. 4**). The effect of larger surface areas at lower altitudes will not much matter at the spatial scale of the home range of frugivores, including megafauna.

Differences in vertical seed dispersal distances among frugivores were not necessarily clear. In the previous studies, mammals with larger home ranges dispersed the seeds of *C. leveilleana* and *A. arguta* farther toward higher or lower altitudes in KAN (*1, 3*). Such a relationship was not detected in the vertical seed dispersal of *C. leveilleana* in ASH. It was surprising that even birds, which are expected to have the smallest home range [except for the jungle crows, whose home range can reach up to 3,000 ha (*21*)], showed strong uphill seed dispersal, as did bears. Efficient use of the steepness around the census route of *C. leveilleana* in ASH by frugivores would affect the results. Altitudes around the census route dramatically change with horizontal distance, from the lowest 900 to 1,400 m a.s.l. at 1,500 m horizontal distances (i.e., a slope of 18°). Such steepness would give frugivores—especially birds which can fly—opportunities to easily visit higher altitudes where different food resources are available because of the altitudinal gradient of the phenological stage (*1*), resulting in strong uphill seed dispersal. Uphill seed dispersal by birds might also be affected by their breeding activity. The timing of *C. leveilleana* fruiting corresponds to the timing of bird breeding (*22*). Birds need to protect nests during the breeding period, while they need to search for many food resources to nurture their chicks. Adherence to their nests while visiting higher altitudes for foods might reinforce uphill seed dispersal. As for *P. grayana* in ASH and ABU, differences in vertical seed dispersal among frugivores and among mountains were not remarkable in terms of both vertical and absolute vertical seed dispersal distance, probably due to the low motivation to ascend or descend mountains. The exception is martens in ABU, which weakly dispersed seeds toward higher altitudes, while birds in ABU and frugivores in ASH weakly dispersed seeds toward lower altitudes. The altitudes of the census route of *P. grayana* in ABU were 550–700 m a.s.l., which are the lowest altitudes compared to those of the other route, correspond to the transitional area between the hilly and montane zones. In the montane zone, *A. arguta*, the fruits of which are the one of the most preferred by mammals in Japan (*3*), are more abundant than in the hilly zone (*23*). We sometimes observed the feces of martens, including seeds of *A. arguta*, when we collected mammal-dispersed seeds of *P. grayana* in the census route. Therefore, the martens in ABU might ascend to visit the montane zones searching for the fruits of *A. arguta*, resulting in the weak uphill seed dispersal of *P. grayana*.

When we combined the amount of seed dispersal and vertical seed dispersal distance, it was evident that bears play a critical role in the vertical seed dispersal of both *C. leveilleana* and *P. grayana* (**Fig. 5**). In both cherry species, bears accounted for ca. 50% of seed dispersal by mammals, and the number of mammal-dispersed seeds in ASH was at most 10 times larger than that in ABU, where bears are locally extinct, despite the fact that the densities of mature cherry trees between ASH and ABU were not much remarkably different. More than 84% of bear-dispersed seeds of *C. leveilleana* in ASH and KAN, and more than 17% of bear-dispersed seeds of *P. grayana* in ASH, were dispersed uphill, and thus contributed to cherry migration toward higher and thus colder altitudes under global warming. Although the effect of bear extinction on the recruitment of *C. leveilleana* and *P. grayana* in terms of the genetic structure of mature trees is not detected at present (*24*), the effect is likely to become clear in the warming world. Macaques dispersed a considerable number of seeds of *C. leveilleana* toward higher altitudes in ASH; however, their contribution to *C. leveilleana* dispersal in KAN was slight and their contribution to *P. grayana* in ASH was not necessarily high. Macaque species, including the Japanese macaque, are omnivores and have very broad food habits. For example, Japanese macaques in a deciduous forest eat woody leaves, flowers, fungi, seeds including nuts, fruits, buds, bark, herbaceous plants, insects, spiders, limpets, and frogs (*25*). Because they choose foods from various candidates in the mountains, both cherry species would not be necessarily chosen despite that they are common fleshy-fruited plants in all mountains. Martens dispersed a considerable number of *C. leveilleana* and *P. grayana* seeds toward higher altitudes in all mountains. It is noticeable that in ABU, where bears and macaques are extinct, martens were apparently the most important frugivorous mammals in both the number of seed dispersal and vertical seed dispersal distance. Other intermediate-sized frugivorous mammals, that is, red foxes, Japanese badgers, and raccoon dogs, which are less frugivory (*12*), contributed little to the vertical seed dispersal of both cherry species in all mountains. Birds contributed to the uphill seed dispersal of both *C. leveilleana* and *P. grayana* in ASH and ABU, although we could not compare their contribution to that of mammals in terms of the number of seed dispersal due to the different sampling methods used. As for *C. leveilleana*, considering that the number of bird-dispersed seeds seems small (**Fig. 2**) and that fruit removal rate by birds is constantly low (ca. 10% for three years) (*7*), the contribution of birds to the overall vertical seed dispersal will be low. As for *P. grayana*, considering that the number of bird-dispersed seeds seems high and that the fruit removal rate by birds can be high (57% at most in the three years studied), the contribution of birds to overall vertical seed dispersal will be high. In areas of Japan where bears are absent, martens and birds that widely inhabit various forests (*26, 27*) may compensate for the loss of bears for both cherry species to some extent. Considering that seed dispersal by marten species and various birds are frequently reported in a wide range of the temperate zones, including North America and Europe (*2, 28*), they are likely to compensate for the lack of vertical seed dispersal of many fleshy-fruited plants by megafauna in many regions where seed dispersal by megafauna is currently not expected. Their compensation may be critically important in Europe (except Russia), where remaining frugivorous megafauna (i.e., brown bears) has almost extinct (*26*).

In this study, we attempted to grasp the whole image of vertical seed dispersal by animals for the first time, and there are several specific caveats to consider in future research. First, there is an inconsistency between vertical and previous horizontal seed dispersal distances. The absolute mean vertical seed dispersal distance of *C. leveilleana* (328.8 m in ASH), and *P. grayana* (105.3 and 172.9 m in ASH and ABU) by birds seems too large compared to the mean horizontal seed dispersal distance of them by birds, which is estimated to be ca. 40–90 m in ABU (*7*). It is possible that the previous study underestimated the horizontal seed dispersal distance; it was estimated based on the assumption that seeds are not dispersed more than 500 m horizontally, in a closed old-growth forest, which failed to evaluate long-distance seed dispersal by jungle crows, which often disperse seeds including cherry ones, outside of closed forests (*29, 30*). In a different model in the same forest, the horizontal seed dispersal distance of *C. leveilleana* was longer than that of the previous one (i.e., more than 50% of seeds were estimated to be dispersed more than 175.7 m) (*31*), suggesting the necessity of validating the estimated horizontal seed dispersal distance. However, combined with the fact that 95% credible intervals of vertical seed dispersal distance are quite large, especially in *C. leveilleana* in ASH (**Fig. 3**), we need to be careful about the validity of mean vertical seed dispersal distance at present, and need to shorten the credible intervals by improving the isotope analysis (*3*). Second, we could not distinguish the bird species of dispersed seeds. Seed dispersal patterns can differ among bird species (*32, 33*). It is necessary to distinguish them to understand the details of vertical seed dispersal by birds using the methods such as the genetic approach (*34*). Third, we conducted the study in forest landscapes. There are open lands, such as pastures, on some human-disturbed mountains, and alpine tundra above forest lines on high mountains. In a mosaic landscape of forests and open lands, forest specialist and habitat generalist frugivores may move—and thus disperse seeds—differently (*2*). The evaluation of vertical seed dispersal in such landscapes, especially in alpine zones, where many endangered plants are distributed, is encouraged. Finally, it is noticeable that we conduct studies in rather midslopes of mountains, not in extreme locations such as the very top of mountains where uphill seed dispersal is not expected, and the very bottom of mountains or plains where downhill seed dispersal is not expected. At such locations, the effect of topography on the direction of vertical seed dispersal may be higher than that of the timing of fruiting.

Our study suggests that the timing of fruiting, rather than topography, determines the direction of vertical seed dispersal by mammals and birds in the temperate zone, irrespective of mountains. In the temperate zone, endozoochorous plants (i.e., fleshy-fruited plants), one of the major types of animal-dispersed plants, account for ca. 35%–42% of woody species in temperate forests (*10*). Synzoochorous plants (i.e., nut plants), another major type of animal-dispersed plants, are dominant in temperate forests in terms of their biomass. Considering that most of these animal-dispersed plants fruit in autumn–winter, our results predict that downhill vertical seed dispersal by animals becomes pervasive and may cause drastic changes in forest composition in the temperate zone. Although we need to investigate plant recruitment after vertical seed dispersal for the judgement of migration success or failure (*4*), there is a concern over the spreading effects of biased vertical seed dispersal by animals on forest ecosystems. In this situation, the role of megafauna in the migration of animal-dispersed plants will be a double-edged sword. We confirmed that megafauna play an essential role in vertical seed dispersal, similar to previous studies investigating horizontal seed dispersal (*14, 16*), although their vertical seed dispersal distance was not necessarily longer than that of other frugivores. Megafauna are expected to be highly helpful to the escape of spring- and summer-fruiting plants under global warming. By contrast, megafauna could be highly harmful to the escape of autumn- and winter-fruiting plants by dispersing many seeds toward lower and thus warmer altitudes. However, it is also true that they disperse a non-negligible number of seeds of autumn-fruiting plants toward higher altitudes (*3*). It is necessary to evaluate whether such uphill dispersal by megafauna effectively facilitates the escape of autumn- and winter-fruiting plants to understand their overall contribution under global warming.

## MATERIAL AND METHODS

### Study site

The study sites were located in the Ashio-Nikko Mountains (ASH, 36.7°N, 139.4–139.5°E), Abukuma Highlands (ABU, 36.9°N, 140.6°E), and Kanto Mountains (KAN, 35.8°N, 139.0°E), central Japan (**Fig. 1A**). The annual mean temperature and precipitation at the nearest weather stations in ASH, ABU, and KAN are 7.2 °C and 2,231 mm (1,292 m a.s.l.), 10.7 °C and 1,910 mm (555 m a.s.l.), and 12.3 °C and 1,610 mm (550 m a.s.l.), respectively (*35, 36*). ASH and KAN comprise steep mountains, and ABU comprises gentle-sloped mountains (**Fig. 1B**). The nearest peaks in ASH, ABU, and KAN around the study sites are Mt. Nantai (2,486 m a.s.l.), Mt. Eizomuro (882 m a.s.l.), and Mt. Hiryu (2,077 m a.s.l.), respectively. The study areas are mountainous and covered with forest vegetation. Natural forests and conifer plantations (*Cryptomeria japonica, Chamaecyparis obtuse*, and, partially, *Larix kaempferi*) cover most of the area (*15, 37, 38*). The natural forest vegetation is similar in the three mountains. In KAN, forests are dominated by *Quercus serrata* and *Castanea crenata* in the hilly zone (0–800 m a.s.l.); *Q. crispula* and *Fagus crenata* in the montane zone (800–1,800 m a.s.l.); and *Abies veitchii* and *Tsuga diversifolia* in the subalpine zone (1,800–2,500 m a.s.l.) (*3*). In ASH and ABU, where the study sites are ca. 100 km north of KAN, while dominant tree species are in common with KAN, the altitude of each vegetation zone is ca. 200 m lower than that in KAN (*15, 38*) (*15, 38*) (Naoe personal observation). ABU lacks the subalpine zone due to its low altitude.

In the three mountains, fauna of frugivorous mammals and birds are relatively well-preserved (*1, 29, 37, 39*–*41*). For example, all of the representative intermediate-sized frugivorous mammals in Japan [red fox (*Vulpes vulpes*), Japanese badger (*Meles anakuma*, previously known as a subspecies of *Meles meles*), raccoon dog (*Nyctereutes procyonoides*), and Japanese marten (*Martes melampus*)] are present. However, larger frugivorous mammals—Asian black bear and Japanese macaque have been locally extinct in ABU for more than decades, probably due to intensive clear-cutting of forests and economic use of remnant forests (*39*). Most frugivorous bird species are observed in both ASH and ABU, where we investigated seed dispersal by birds, but the composition of dominant birds differs between the mountains. In ASH, the dominant frugivorous birds are the narcissus flycatcher (*Ficedula narcissina*), brown-headed thrush (*Turdus chrysolaus*), Asian brown flycatcher (*Muscicapa dauurica*), and Eurasian jay (*Garrulus glandarius*) (*40*). In ABU, the dominant frugivorous birds are the brown-eared bulbul (*Hypsipetes amaurotis*), narcissus flycatcher (*F. narcissina*), grey thrush (*T. cardis*), and Japanese white-eye (*Zosterops japonicus*) (*22*). The largest frugivorous bird in Japan (*29*), the jungle crow (*Corvus macrorhynchos*), is the fifth-most dominant in both mountains.

### Plant materials

We targeted two fleshy-fruited cherry species common in mountainous areas in central Japan: the wild flowering cherry *Cerasus leveilleana* (Koehne) H. Ohba (syn. *Prunus verecunda*) and the Japanese bird cherry *Padus grayana* (Maxim.) C. K. Schneid (syn. *Prunus grayana*). *Cerasus leveilleana* is a deciduous canopy tree species with a minimum reproductive diameter at breast height (DBH) of 11.8 cm (*22*). It produces ball-shaped red to black fleshy fruits (7.0 mm in diameter) from mid-June to mid-July (*17*). *Padus grayana* is a deciduous subcanopy tree species with a minimum reproductive DBH of 7.2 cm. It produces tear-shaped red to black fleshy fruits (7.0 mm and 5.2 mm in longer and shorter diameter) from August to mid-September. Their fruits are dispersed by various mammals and birds (*9, 17, 42*).

### Sampling of dispersed and non-dispersed seeds

Samplings of mammal- and bird-dispersed seeds were conducted in 2014 and 2015 for *C. leveilleana*, and in 2014 and 2016 for *P. grayana*, when the fruit production of each cherry species was high. To collect dispersed seeds, we set two 2 km-census routes in both ASH (850– 1,200 m a.s.l.) and ABU (550–750 m a.s.l.) along paved roads mainly going through natural forests (3–4 m in width) (for details, see **Table S1**). One route was set for *C. leveilleana*, and another was set for *P. grayana*, where the target cherry species were abundant. The densities of mature trees along and around the routes in ASH and ABU were 13.1 and 13.0 individuals per km for *C. leveilleana*, and 22.3 and 11.5 per km for *P. grayana*, respectively (*24*). To collect mammal-dispersed seeds, we visited the routes weekly during the fruiting season of each cherry species, and examined all fresh mammalian feces found along the routes in situ for the seeds of the cherry species. Mammalian species were identified based on the shape and smell of the feces (*37*). Each fecal sample was washed through a series of sieves, after which the number of mature intact seeds was counted. To collect bird-dispersed seeds, we set 40 seed traps along the routes in both ASH and ABU. Seed traps had a surface area of 0.5 m^2^ and were made of nylon cloth (mesh size of 1 mm). The seed traps were set 1 m above the ground to avoid predation by woody mice. When we set seed traps for the first time (for *C. leveilleana* in 2014), 20 seed traps were set rather regularly, and 20 traps were set just outside the tree crown of the fruiting target cherry species. But this method failed to sample sufficient bird-dispersed seeds. Thus, we set 20 traps rather regularly and 20 traps under the tree crown of them in the subsequent samplings (i.e., for *C. leveilleana* in 2015 and *P. grayana* in 2014 and 2016). We collected seeds that fell into the traps every two weeks during the fruiting seasons. Pulpless seeds were considered to have been dispersed by birds through either regurgitation or defecation. The locations of all sampling points were recorded using handheld GPS receivers (GPSMAP 60CSx, 62S, or Oregon 550; Garmin Ltd., Olathe, KS, USA).

We collected non-dispersed reference seeds from fruiting trees at various altitudes. This is because it is necessary to confirm the negative correlation between altitudes and the oxygen isotope ratio of non-dispersed seeds, and use it as the calibration line to estimate the vertical seed dispersal distance (*18*). This sampling was conducted for *C. leveilleana* in ASH in 2015 and for *P. grayana* in ASH and ABU in 2016. For seed sampling details in KAN, please see the literature (*1*). Briefly, mammal-dispersed seeds of *C. leveilleana* were collected along and around a 16 km-census route (550–1,650 m a.s.l.) at 10-day intervals during the fruiting season in 2010–2013. The subsequent methods were the same as those conducted in ASH and ABU.

### Estimation of vertical seed dispersal distance

The estimation of vertical seed dispersal distance was conducted based on the method described in the literature, as follows (*3*): first, we confirmed the negative correlation between altitudes and the oxygen isotope ratio of non-dispersed seeds; second, we estimated the altitudes of the mother trees of dispersed seeds by using the negative correlation (i.e., by using the regression equation as a calibration) and the oxygen isotope ratio of dispersed seeds; third, we calculated the vertical seed dispersal distance by subtracting the altitudes of the mother trees from those of the dispersed seeds.

Calibration lines can vary annually owing to climatic factors (*18*). This variation may cause uncertain errors in vertical seed dispersal estimates if the sampling year of dispersed seeds is different from that of non-dispersed seeds for calibration. Considering this carefully, we only used dispersed seeds that were collected in the same year that the calibration line was made for estimation (*C. leveilleana*, seeds collected in ASH in 2015; *P. grayana*, seeds collected in ASH in 2016 and ABU in 2016). For KAN, dispersed seeds were collected in 2010–2013 and non-dispersed seeds in 2012 and 2013 (*1*). Following the previous study (*1*), we applied the calibration line in 2012 to estimate the vertical seed dispersal distance of dispersed seeds collected in 2010 and 2011. This is because the correlation is extremely strong between the oxygen isotope ratios of non-dispersed seeds and mean air temperature in May in KAN, when the seeds are developing (*r* = 0.9487, *n* = 15, *P* < 0.0001), and because the air temperatures in May 2010–2012 were almost identical (14.9–15.1°C) (*1*). We did not conduct this procedure for the dispersed seeds in ASH and ABU, because differences in air temperatures among the sampling years of dispersed and non-dispersed seeds were relatively large (> 0.5°C) in most cases (*3*), and because the relationship between the oxygen isotope ratios of non-dispersed seeds and mean air temperature was not examined in ASH and ABU.

We measured the oxygen isotope ratio of the endocarps of the seeds, which are the most external dead and hard tissue of the diaspores of cherry species (*18*). For non-dispersed seeds to make calibration lines, we analyzed three seeds at each altitude (*1*). For dispersed seeds, we randomly selected one seed from each fecal or seed-trap sample and analyzed them. We measured the oxygen isotope ratio by continuous flow (CF) combustion and isotope ratio mass spectrometry (IRMS) analysis using a high-temperature elemental analyzer (TC/EA; Thermo Fisher Scientific, Waltham, MA, USA) coupled online with a mass spectrometer (Delta Plus XP; Thermo Fisher Scientific) using a ConFlo III interface (Thermo Fisher Scientific). Detailed methods are described in the literature (*18*). The sample gases were calibrated by measuring the reference substances (IAEA-601 and IAEA-602 benzoic acid; 23.14‰ and 71.28‰, respectively) of known isotope compositions (*43*). The isotopic composition of a sample is conventionally expressed as the ‘δ’ value by comparison with the international primary reference material (V-SMOW) as follows:

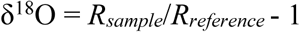

where *R* denotes the ratio of numbers (*N*) of each isotope, as follows:

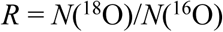

The standard deviation of the replicates was approximately 0.2‰ (1*σ*) for the measurements.

### Evaluating the effect of topography on vertical seed dispersal

To examine whether the topography of the mountains affect vertical seed dispersal, we estimated the expected vertical seed dispersal by simulation, assuming that frugivores randomly move and disperse seeds based on the topography in their home ranges. We used the locations where dispersed seeds were collected and topography around the locations for the simulation. One location was randomly sampled from a circle with a horizontal radius of *r* km centered on each location where dispersed seeds were collected. To sufficiently reflect the topography, we conducted this procedure five times per dispersed seed. Then, the altitudinal distribution was obtained by summarizing these locations. A circle assumes the home range of frugivores. We tried different values for *r*: 0.1, 0.5, 1, 3, and 10 km, which result in 3, 75, 300, 2,700, and 30,000 ha in the circular area, respectively. These are roughly correspondent to the home range sizes of small birds, intermediate-sized birds, large birds and intermediate-size mammal, Asian black bears in KAN, and Asian black bears in ASH, respectively (*26, 27*). These simulations were performed for each cherry species and site.

### Contribution of each frugivore to whole vertical seed dispersal

To assess the contribution of each frugivore to whole vertical seed dispersal, we multiplied the number of dispersed seeds and vertical seed dispersal distance of each frugivore, and generated histograms for each cherry species and mountains. Because we only analyzed one dispersed seed per feces because of the labor involved, we assumed that the vertical seed dispersal distance of remaining the non-analyzed seeds in each feces was identical to that of analyzed seed in the same feces.

### Statistical analysis

When we estimated the vertical seed dispersal distance based on the calibration lines, Bayesian inference was used to evaluate errors in the altitude identification of the mother tree, and thus, the vertical seed dispersal distance. Uniform distributions were used as the prior distributions for all parameters. We ran the Markov chain Monte Carlo (MCMC) algorithm for three independent chains of 110,000 iterations. In each chain, the first 10,000 iterations were abandoned as a burn-in and the remaining chain was thinned every 50 steps, resulting in 2,000 values per chain sampled from the posterior. Finally, 6,000 samples were obtained from three chains, which were used to yield the posterior distributions and summarize the parameters. Model convergence was assessed visually and using the *R^* values (the Gelman–Rubin statistic) (*19*). These analyses were performed using R version 3.5.2 (*44*) and WinBUGS version 1.4.3 (*45*).

## Supplementary Materials

Fig. S1. Calibration lines (i.e., negative correlation between altitudes and the isotope ratio of non-dispersed reference seeds) for the vertical seed dispersal distance of (**A**) *Cerasus leveilleana* and **(B)** *Padus grayana*. Dots indicate non-dispersed reference seeds, and lines indicate calibration lines.

Table S1. Detailed sampling of the target species *Cerasus leveilleana* and *Padus grayana* in the Ashio-Nikko Mountains (ASH), Abukuma Highlands (ABU), and Kanto Mountains (KAN). We only estimated the vertical seed dispersal distance of dispersed seeds that were collected in the same year that the calibration line was made (see Methods for details).

Table S2. Number of dispersed and analyzed seeds of *Cerasus leveilleana* and *Padus grayana*.

Table S3. Results of Bayesian inference for calibration lines (oxygen isotope ratio of seed = *a* × altitude + *b*) of *Cerasus leveilleana* and *Padus grayana*.

Table S4. Observed and expected absolute mean vertical seed dispersal distances of *Cerasus leveilleana* and *Padus grayana* in ASH, ABU, and KAN.

## ACKNOWLEDGEMETNS

**General**: We thank Tadashi Iwasaki, Hiroaki Myoujou and Tetsuro Yoshikawa for their field assistance in seed sampling; Rie Takaya and Chikage Yoshimizu for their laboratory assistance, and Mitsue Shibata for their comments. **Funding:** S.N., Si.K., I.T., Te.N., Sh.K., Sa.K., and T.M. were funded by JSPS KAKENHI Grant Numbers 25241026, 15K18718, 16H02524, 17H00797, and 17H05031. This study was conducted by the support of Joint Research Grant for the Environmental Isotope Study of Research Institute for Humanity and Nature. **Author contributions:** S.N. conceived the idea; S.N., Si.K., Te.N., and T.M. contributed to the study design; S.N., Si.K., and To.N. collected samples; S.N., Y.T., Sh.K., Sa.K., I.T., and T.H. conducted laboratory work; Y.T. analyzed the data; S.N. and Y.T. wrote the manuscript with input from all authors. **Competing interests:** The authors declare that they have no competing interests. **Data and materials availability:** All data needed to evaluate the conclusions in the paper are present in the paper and/or the Supplementary Materials.

## REFERENCES

1. S. Naoe, I. Tayasu, Y. Sakai, T. Masaki, K. Kobayashi, A. Nakajima, Y. Sato, K. Yamazaki, H. Kiyokawa, S. Koike, Mountain-climbing bears protect cherry species from global warming through vertical seed dispersal. Curr. Biol. 26, R315–R316 (2016).

2. J. P. González-Varo, J. V. López-Bao, J. Guitián, Seed dispersers help plants to escape global warming. Oikos 126, 1600–1606 (2017).

3. S. Naoe, I. Tayasu, Y. Sakai, T. Masaki, K. Kobayashi, A. Nakajima, Y. Sato, K. Yamazaki, H. Kiyokawa, S. Koike, Downhill seed dispersal by temperate mammals: a potential threat to plant escape from global warming. Sci. Rep. 9, 14932 (2019).

4. R. T. Corlett, D. A. Westcott, Will plant movements keep up with climate change? Trends Ecol. Evol. 28, 482–488 (2013).

5. R. G. Barry, R. J. Chorley, Atmosphere, Weather and Climate (Routledge, London, ed. 9, 2009).

6. R. Cousens, C. Dytham, R. Law, Dispersal in Plants: A Population Perspective (Oxford University Press, Oxford, 2008).

7. S. Naoe, T. Masaki, S. Sakai, Effects of temporal variation in community-level fruit abundance on seed dispersal by birds across woody species. Am. J. Bot. 105, 1792–1801 (2018).

8. T. Yoshikawa, S. Harasawa, Y. Isagi, N. Niikura, S. Koike, H. Taki, S. Naoe, T. Masaki, Relative importance of landscape features, stand structural attributes, and fruit availability on fruit-eating birds in Japanese forests fragmented by coniferous plantations. Biol. Conserv. 209, 356–365 (2017).

9. T. Yoshikawa, Y. Isagi, K. Kikuzawa, Relationships between bird-dispersed plants and avian fruit consumers with different feeding strategies in Japan. Ecol. Res. 24, 1301–1311 (2009).

10. C. M. Herrera, in Plant–animal interactions: an evolutionary approach, C. M. Herrera, O. Pellmyr, Eds. (Blackwell Publishing, Oxford, 2002), pp. 185–208.

11. T. Rötzer, F. M. Chmielewski, Phenological maps of Europe. Clim. Res. 18, 249–257 (2001).

12. S. Koike, S. Kasai, K. Yamazaki, K. Furubayashi, Fruit phenology of *Prunus jamasakura* and the feeding habit of the Asiatic black bear as a seed disperser. Ecol. Res. 23, 385–392 (2008).

13. C. Körner, E. M. Spehn, in Mountain Biodiversity: A Global Assessment, C. Korne, E. M. Spehn, Eds. (Parthenon Publishing, New York, 2002), pp. 3–20.

14. A. Campos-Arceiz, S. Blake, Megagardeners of the forest - the role of elephants in seed dispersal. Acta Oecol. 37, 542–553 (2011).

15. S. Koike, T. Masaki, Y. Nemoto, C. Kozakai, K. Yamazaki, S. Kasai, A. Nakajima, K. Kaji, Estimate of the seed shadow created by the Asiatic black bear *Ursus thibetanus* and its characteristics as a seed disperser in Japanese cool-temperate forest. Oikos 120, 280–290 (2011).

16. K. R. McConkey, A. Nathalang, W. Y. Brockelman, C. Saralamba, J. Santon, U. Matmoon, R. Somnuk, K. Srinoppawan, Different megafauna vary in their seed dispersal effectiveness of the megafaunal fruit *Platymitra macrocarpa* (Annonaceae). PLoS One 13, 1–18 (2018).

17. T. Masaki, K. Takahashi, A. Sawa, T. Kado, S. Naoe, S. Koike, M. Shibata, Fleshy fruit characteristics in a temperate deciduous forest of Japan: how unique are they? J. Plant Res. 125, 103–114 (2012).

18. S. Naoe, I. Tayasu, T. Masaki, S. Koike, Negative correlation between altitudes and oxygen isotope ratios of seeds: exploring its applicability to assess vertical seed dispersal. Ecol. Evol. 6, 6817–6823 (2016).

19. A. Gelman, J. Hill, Data Analysis Using Regression and Multilevel/Hierarchical Models (Cambridge University Press, Cambridge, 2007).

20. C. Kozakai, K. Yamazaki, Y. Nemoto, A. Nakajima, Y. Umemura, S. Koike, Y. Goto, S. Kasai, S. Abe, T. Masaki, K. Kaji, Fluctuation of daily activity time budgets of Japanese black bears: relationship to sex, reproductive status, and hard-mast availability. J. Mammal. 94, 351–360 (2013).

21. N. Fujita, T. Hattori, A. Azuma, O. Mai, M. Yazawa, N. Segawa, Determining Characteristics of Jungle Crow’s Home-range and a Planning Area for Developing a Policy to Control Crows Population. J. Rural Plan. Assoc. 34, 160–166 (2015).(in Japanese with English summary)

22. S. Naoe, S. Sakai, A. Sawa, T. Masaki, Seasonal difference in the effects of fragmentation on seed dispersal by birds in Japanese temperate forests. Ecol. Res. 26, 301–309 (2011).

23. Y. Horikawa, Atlas of the Japanese Flora: An Introduction to Plant Sociology of East Asia (Gakken, Tokyo, 1972).

24. T. Nagamitsu, K. Shuri, S. Kikuchi, S. Koike, S. Naoe, T. Masaki, Multiscale spatial genetic structure within and between populations of wild cherry trees in nuclear genotypes and chloroplast haplotypes. Ecol. Evol. 9, 11266–11276 (2019).

25. Y. Tsuji, S. Fujita, H. Sugiura, C. Saito, S. Takatsuki, Long-term variation in fruiting and the food habits of wild Japanese macaques on Kinkazan Island, northern Japan. Am. J. Primatol. 68, 1068–1080 (2006).

26. S. D. Ohdachi, Y. Ishibashi, M. A. Iwasa, T. Saitioh, The Wild Mammals of Japan (Shoukadoh, Kyoto, ed. 2, 2015).

27. H. Higuchi, H. Morioka, S. Yamagishi, The Encyclopaedia of Animals in Japan, Volume 4: Birds 2 (Heibonsya Limited, Tokyo, 1997), vol. 4. (in Japanese)

28. B. Snow, D. W. Snow, Birds and Berries: A Study of an Ecological Interaction (T & AD Poyser, London, 1988).

29. S. Naoe, My forest study: focusing on seed dispersal by birds (Saela Shobo, Tokyo, 2015). (in Japanese)

30. H. Nishi, S. Tsuyuzaki, Seed dispersal and seedling establishment of *Rhus trichocarpa* promoted by a crow (*Corvus macrorhynchos*) on a volcano in Japan. Ecography 3, 311–323 (2004).

31. T. Masaki, T. Nakashizuka, K. Niiyama, H. Tanaka, S. Iida, J. M. Bullock, S. Naoe, Impact of the spatial uncertainty of seed dispersal on tree colonization dynamics in a temperate forest. Oikos 128, 1816–1828 (2019).

32. P. Jordano, C. García, J. A. Godoy, J. L. García-Castaño, Differential contribution of frugivores to complex seed dispersal patterns. Proc. Natl. Acad. Sci. U. S. A. 104, 3278–82 (2007).

33. Y. Tsunamoto, S. Naoe, T. Masaki, Y. Isagi, Different contributions of birds and mammals to seed dispersal of a fleshy-fruited tree. Basic Appl. Ecol. 43, 66–75 (2020).

34. J. P. González-Varo, J. M. Arroyo, P. Jordano, Who dispersed the seeds? The use of DNA barcoding in frugivory and seed dispersal studies. Methods Ecol. Evol. 5, 806–814 (2014).

35. Y. Moriguchi, T. Morishita, Y. Ohtani, in Diversity and interaction in a temperate forest community: Ogawa Forest Reserve of Japan, T. Nakashizuka, Y. Matsumoto, Eds. (Springer, Tokyo, 2002), pp. 11–18.

36. Japan Meteorological Agency, Mean monthly meteorological data at Ogouchi between 1981–2010 (2016) (available at http://www.data.jma.go.jp/obd/stats/etrn/view/nml_amd_ym.php?prec_no=44&block_no=0365&year=&month=&day=&view=p1). (in Japanese)

37. S. Koike, H. Morimoto, Y. Goto, C. Kozakai, K. Yamazaki, Frugivory of carnivores and seed dispersal of fleshy fruits in cool-temperate deciduous forests. J. For. Res. 13, 215–222 (2008).

38. Y. Yamaura, S. Ikeno, M. Sano, K. Okabe, K. Ozaki, Bird responses to broad-leaved forest patch area in a plantation landscape across seasons. Biol. Conserv. 142, 2155–2165 (2009).

39. K. Yamazaki, K. Koyanagi, A. Tsuji, List of Mammals Found in Ibraki Prefecture, Central Part of Japan. Bull. Ibaraki Nat. Museum. 4, 103–108 (2001). (in Japanese with English summary)

40. K. Okuda, Y. Seki, M. Koganezawa, Changes in the community structure of breeding birds in Oku-Nikko, Japan, with special reference to increases in the sika deer population. Japanese J. Conserv. Ecol. 76, 69–76 (2013). (in Japanese with English summary)

41. A. Inagaki, M. L. Allen, T. Maruyama, K. Yamazaki, K. Tochigi, T. Naganuma, S. Koike, Vertebrate scavenger guild composition and utilization of carrion in an East Asian temperate forest. Ecol. Evol. 10, 1223–1232 (2020).

42. S. Koike, T. Masaki, Characteristics of fruits consumed by mammalian frugivores in Japanese temperate forest. Ecol. Res. 34, 246–254 (2019).

43. W. A. Brand, T. B. Coplen, A. T. Aerts-Bijma, J. K. Böhlke, M. Gehre, H. Geilmann, M. Gröning, H. G. Jansen, H. A. J. Meijer, S. J. Mroczkowski, H. Qi, K. Soergel, H. Stuart-Williams, S. M. Weise, R. A. Werner, Comprehensive inter-laboratory calibration of reference materials for d18O versus VSMOW using various on-line high-temperature conversion techniques. Rapid Commun. Mass Spectrom. 23, 999–1019 (2009).

44. R Development Core Team, R: A language and environment for statistical computing (R Foundation for Statistical Computing, Vienna, 2018; http://www.r-project.org).

45. D. J. Lunn, A. Thomas, N. Best, D. Spiegelhalter, WinBUGS –A Bayesian modelling framework: Concepts, structure, and extensibility. Stat. Comput. 10, 325–337 (2000).

