## Supplementary material for "Timing of fruiting rather than topography determines the direction of vertical seed dispersal by mammals and birds": Table S1

### Supplementary Materials

Fig. S1. Calibration lines (i.e., negative correlation between altitudes and the isotope ratio of non-dispersed reference seeds) for the vertical seed dispersal distance of (**A**) *Cerasus leveilleana* and (**B**) *Padus grayana*. Dots indicate non-dispersed reference seeds, and lines indicate calibration lines.

Table S1. Detailed sampling of the target species *Cerasus leveilleana* and *Padus grayana* in the Ashio-Nikko Mountains (ASH), Abukuma Highlands (ABU), and Kanto Mountains (KAN). We only estimated the vertical seed dispersal distance of dispersed seeds that were collected in the same year that the calibration line was made (see Methods for details).

Table S2. Number of dispersed and analyzed seeds of *Cerasus leveilleana* and *Padus grayana*.

Table S3. Results of Bayesian inference for calibration lines (oxygen isotope ratio of seed = $a \times$ altitude + $b$) of *Cerasus leveilleana* and *Padus grayana*.

Table S4. Observed and expected absolute mean vertical seed dispersal distances of *Cerasus leveilleana* and *Padus grayana* in ASH, ABU, and KAN.

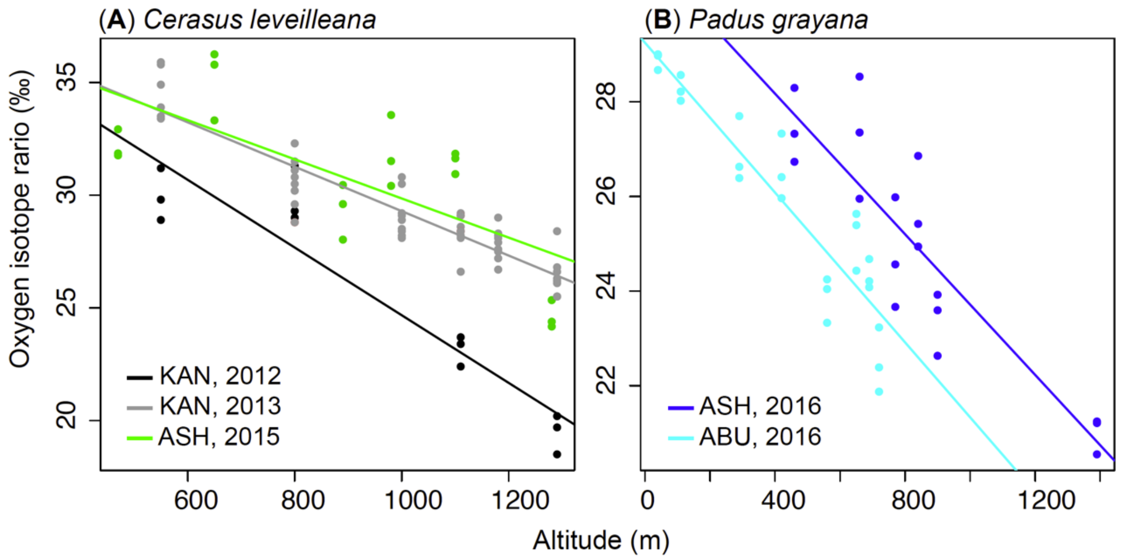


Fig. S1. Calibration lines (i.e., negative correlation between altitudes and the isotope ratio of non-dispersed reference seeds) for the vertical seed dispersal distance of (**A**) *Cerasus leveilleana* and (**B**) *Padus grayana*. Dots indicate non-dispersed reference seeds, and lines indicate calibration lines.

Table S1. Detailed sampling of the target species *Cerasus leveilleana* and *Padus grayana* in the Ashio-Nikko Mountains (ASH), Abukuma Highlands (ABU), and Kanto Mountains (KAN). We only estimated the vertical seed dispersal distance of dispersed seeds which were collected in the same year that the calibration line was made (see Methods for details).

| Mountains | Latitude | Longitude | Species | Vertical distribution of species (m) | Census altitude (m) | Census route (km) | Target frugivore | Sampling year | Seed dispersal quantity | Vertical seed dispersal distance |
|---|---|---|---|---|---|---|---|---|---|---|
| ASH | 36.7°N | 139.5°E | *C. leveilleana* | 450–1,150m | 1,050–1,200m | 2 | mammal, bird | 2014 | ✓† | |
| | | | | | | | | 2015 | ✓† | ✓ |
| | | | *P. grayana* | 450–1,400m | 850–950m | 2 | mammal, bird | 2014 | ✓ | |
| | | | | | | | | 2016 | ✓ | ✓ |
| ABU | 36.9°N | 140.6°E | *C. leveilleana* | 400–880m(summit) | 600–750m | 2 | mammal, bird | 2014 | ✓† | |
| | | | | | | | | 2015 | ✓† | ✓ |
| | | | *P. grayana* | 0–880m(summit) | 550–700m | 2 | mammal, bird | 2014 | ✓ | |
| | | | | | | | | 2016 | ✓ | ✓ |
| KAN* | 35.8°N | 139.0°E | *C. leveilleana* | 550–1,300m | 550–1,650m | 16 | mammal | 2010-2013 | ✓ | ✓ |

*Data are from Naoe et al. 2016.

†Sampling method of bird-dispersed seed between years are different. See Methods for details.

Table S2. Number of dispersed and analyzed seeds of *Cerasus leveilleana* and *Padus grayana*.

| Species | Mountains | Year | Frugivore | No. of dispersed seeds | No. of analyzed seeds |
|---|---|---|---|---|---|
| *C. leveilleana* | ASH | 2014 | Bear | 850 | 0 |
| | | | Macaque | 813 | 0 |
| | | | Raccoon dog | 99 | 0 |
| | | | Marten | 175 | 0 |
| | | | Bird | 29 | 0 |
| | ABU | 2014 | Raccoon dog | 147 | 0 |
| | | | Marten | 577 | 0 |
| | | | Bird | 8 | 0 |
| | ASH | 2015 | Bear | 3029 | 13 |
| | | | Macaque | 770 | 43 |
| | | | Marten | 1348 | 66 |
| | | | Bird | 105 | 23 |
| | ABU | 2015 | Marten | 513 | 0 |
| | | | Bird | 14 | 0 |
| *P. grayana* | ASH | 2014 | Bear | 1688 | 0 |
| | | | Macaque | 733 | 0 |
| | | | Marten | 723 | 0 |
| | | | Bird | 73 | 0 |
| | ABU | 2014 | Marten | 1175 | 0 |
| | | | Raccoon dog | 40 | 0 |
| | | | Bird | 1509 | 0 |
| | ASH | 2016 | Bear | 3066 | 17 |
| | | | Macaque | 32 | 1 |
| | | | Marten | 1495 | 28 |
| | | | Bird | 170 | 47 |
| | ABU | 2016 | Raccoon dog | 240 | 2 |
| | | | Marten | 595 | 18 |
| | | | Bird | 121 | 26 |
| Total | | | | 20137 | 284 |

Table S3. Results of Bayesian inference for calibration lines (oxygen isotope ratio of seed = $a \times$ altitude + $b$) of *Cerasus leveilleana* and *Padus grayana*.

| Species | Mountains | Year | Parameter | Mean | SD | 2.5% | 25% | 50% | 75% | 97.5% | Rhat |
|---|---|---|---|---|---|---|---|---|---|---|---|
| *C. leveilleana* | ASH | 2015 | a | -0.0087 | 0.0025 | -0.0136 | -0.0103 | -0.0087 | -0.0071 | -0.0039 | 1.0016 |
| | | | b | 38.5 | 2.3 | 34.0 | 37.0 | 38.5 | 40.0 | 43.2 | 1.0017 |
| | KAN | 2012 | a | -0.0150 | 0.0021 | -0.0193 | -0.0164 | -0.0151 | -0.0137 | -0.0106 | 1.0011 |
| | | | b | 39.7 | 2.1 | 35.4 | 38.4 | 39.7 | 41.0 | 43.9 | 1.0011 |
| | | 2013 | a | -0.0099 | 0.0007 | -0.0111 | -0.0103 | -0.0099 | -0.0094 | -0.0085 | 1.0008 |
| | | | b | 39.2 | 0.7 | 37.8 | 38.7 | 39.2 | 39.6 | 40.5 | 1.0009 |
| *P. grayana* | ASH | 2016 | a | -0.0074 | 0.0011 | -0.0095 | -0.0081 | -0.0074 | -0.0067 | -0.0053 | 1.0009 |
| | | | b | 31.1 | 0.9 | 29.3 | 30.6 | 31.2 | 31.7 | 33.0 | 1.0008 |
| | ABU | 2016 | a | -0.0079 | 0.0007 | -0.0094 | -0.0084 | -0.0079 | -0.0074 | -0.0064 | 1.0009 |
| | | | b | 29.2 | 0.4 | 28.5 | 29.0 | 29.3 | 29.5 | 30.0 | 1.0008 |

Table S4. Observed and expected absolute mean vertical seed dispersal distances of *Cerasus leveilleana* and *Padus grayana* in the Ashio-Nikko Mountains (ASH), Abukuma Highlands (ABU), and Kanto Mountains (KAN).

| Species | Mountains | Observed absolute mean vertical seed dispersal distance (m) | | | | | Expected absolute mean vertical seed dispersal distance (m) | | | | |
|---|---|---|---|---|---|---|---|---|---|---|---|
| | | Bear | Macaque | Raccoon dog | Marten | Bird | $r = 0.1$ | $r = 0.5$ | $r = 1$ | $r = 3$ | $r = 10$ |
| *C. leveilleana* | ASH | 292.9 ± 29.3 | 314.6 ± 20.9 | | 333.6 ± 18.6 | 328.8 ± 26.6 | 13.2 ± 0.4 | 46.8 ± 1.5 | 67.2 ± 2.1 | 137.8 ± 4.2 | 237.2 ± 7.4 |
| | KAN | 330.6 ± 28.8 | | | 235.1 ± 23.1 | | 27.3 ± 1.1 | 81.4 ± 3.1 | 146.2 ± 5.2 | 231.4 ± 7.3 | 302.9 ± 9.3 |
| *P. grayana* | ASH | 122.5 ± 2.3 | 31.3 | | 132.2 ± 23.0 | 105.3 ± 12.2 | 21.4 ± 0.9 | 59.0 ± 2.4 | 104.9 ± 4.0 | 196.3 ± 7.6 | 318.4 ± 11.6 |
| | ABU | | | 56.5 ± 19.3 | 120.0 ± 18.8 | 172.9 ± 21.6 | 8.7 ± 0.5 | 24.8 ± 1.5 | 42.5 ± 2.5 | 75.3 ± 3.4 | 86.5 ± 4.7 |
